# A mathematical model for simulating the transmission of Wuhan novel Coronavirus

**DOI:** 10.1101/2020.01.19.911669

**Authors:** Tianmu Chen, Jia Rui, Qiupeng Wang, Zeyu Zhao, Jing-An Cui, Ling Yin

**Author notes:** Correspondence: Tianmu Chen, State Key Laboratory of Molecular Vaccinology and Molecular Diagnostics, School of Public Health, Xiamen University, 4221-117 South Xiang’an Road, Xiang’an District, Xiamen, Fujian Province, People’s Republic of China.

## Abstract

As reported by the World Health Organization, a novel coronavirus (2019-nCoV) was identified as the causative virus of Wuhan pneumonia of unknown etiology by Chinese authorities on 7 January, 2020. In this study, we developed a Bats-Hosts-Reservoir-People transmission network model for simulating the potential transmission from the infection source (probable be bats) to the human infection. Since the Bats-Hosts-Reservoir network was hard to explore clearly and public concerns were focusing on the transmission from a seafood market (reservoir) to people, we simplified the model as Reservoir-People transmission network model. The basic reproduction number (*R*_0_) was calculated from the RP model to assess the transmissibility of the 2019-nCoV.

## Introduction

On 31 December 2019, the World Health Organization (WHO) China Country Office was informed of cases of pneumonia of unknown etiology (unknown cause) detected in Wuhan City, Hubei Province of China, and WHO reported that a novel coronavirus (2019-nCoV) was identified as the causative virus by Chinese authorities on 7 January(*1*). Potential for international spread via commercial air travel had been assessed(*2*). Public health concerns have been paid globally on how many people had been infected actually.

In this study, we developed a Bats-Hosts-Reservoir-People (BHRP) transmission network model for simulating the potential transmission from the infection source (probable be bats) to the human infection. Since the Bats-Hosts-Reservoir network was hard to explore clearly and public concerns were focusing on the transmission from a seafood market (reservoir) to people, we simplified the model as Reservoir-People (RP) transmission network model. The basic reproduction number (*R*_0_) was calculated from the RP model to assess the transmissibility of the 2019-nCoV.

### The Bats-Hosts-Reservoir-People transmission network model

We assumed that the virus transmitted among the bats population, and then transmitted to an unknown host (probably be wild animals). The hosts were hunted and sent to the seafood market which was defined as the reservoir or the virus. People exposed to the market got the risks of the infection (Figure 1).

**Figure 1.**
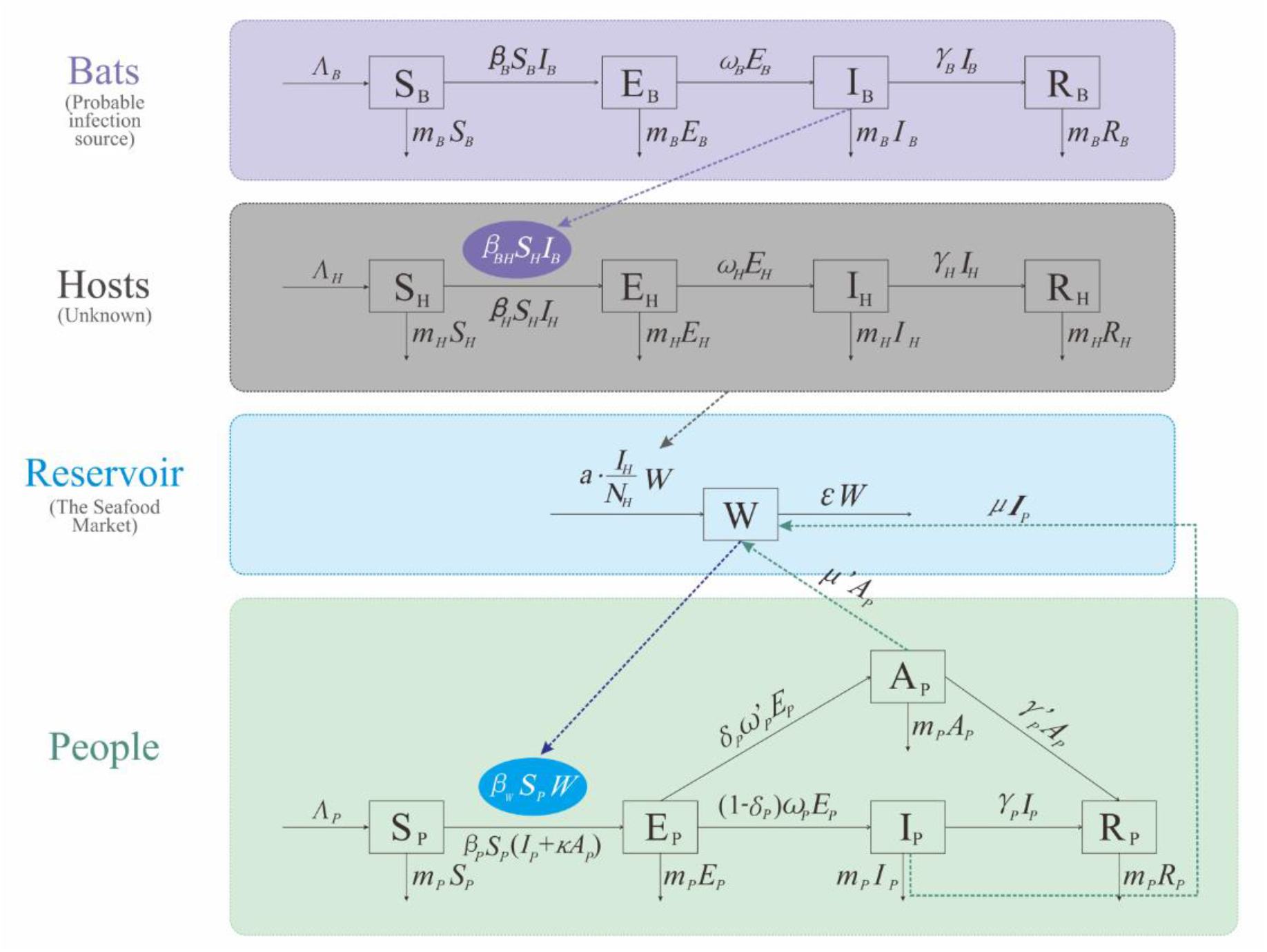
Flowchart of the Bats-Hosts-Reservoir-People transmission network model.

The BHRP transmission network model was based on the following assumptions or facts:

a. The bats were divided into four departments: susceptible bats (*S*_*B*_), exposed bats (*E*_*B*_), infected bats (*I*_*B*_), and removed bats (*R*_*B*_). The birth rate and death rate of bats were defined as *n*_*B*_ and *m*_*B*_. In this model, we set *Λ*_*B*_ *= n*_*B*_ × *N*_*B*_ where *N*_*B*_ refer to the total number of bats. The incubation period of bat infection was defined as 1/*ω*_*B*_ and the infectious period of bat infection was defined as 1/*γ*_*B*_. The *S*_*B*_ will be infected through sufficient contact with *I*_*B*_, and the transmission rate was defined as *β*_*B*_.
b. The hosts were divided into four departments: susceptible hosts (*S*_*H*_), exposed hosts (*E*_*H*_), infected hosts (*I*_*H*_), and removed hosts (*R*_*H*_). The birth rate and death rate of hosts were defined as *n*_*H*_ and *m*_*H*_. In this model, we set *Λ*_*H*_ *= n*_*H*_ × *N*_*H*_ where *N*_*H*_ refer to the total number of hosts. The incubation period of host infection was defined as 1/*ω*_*H*_ and the infectious period of host infection was defined as 1/*γ*_*H*_. The *S*_*H*_ will be infected through sufficient contact with *I*_*B*_ and *I*_*H*_, and the transmission rates were defined as *β*_*BH*_ and *β*_*H*_, respectively.
c. The 2019-nCoV in reservoir (the seafood market) was denoted as *W*. We assumed that the retail purchases rate of the hosts in the market was *a*, and that the prevalence of 2019-nCoV in the purchases was *I*_*H*_/*N*_*H*_, therefore, the rate of the 2019-nCoV in *W* imported form the hosts was *aWI*_*H*_/*N*_*H*_ where *N*_*H*_ was the total number of hosts. We also assumed that symptomatic infected people and asymptomatic infected people could export the virus into *W* with the rate of *μ*_*P*_ and *μ’*_*P*_, although this assumption might occur in a low probability. The virus in *W* will subsequently leave the *W* compartment at a rate of *εW*, where 1/*ε* is the lifetime of the virus.
d. The people were divided into five departments: susceptible people (*S*_*P*_), exposed people (*E*_*P*_), symptomatic infected people (*I*_*P*_), asymptomatic infected people (*A*_*P*_), and removed people (*R*_*P*_) including recovered and death people. The birth rate and death rate of people were defined as *n*_*P*_ and *m*_*P*_. In this model, we set *Λ*_*P*_ *= n*_*P*_ × *N*_*P*_ where *N*_*P*_ refer to the total number of people. The incubation period and latent period of human infection was defined as 1/*ω*_*P*_ and 1/*ω’* _*P*_. The infectious period of *I*_*P*_ and *A*_*P*_ was defined as 1/*γ*_*P*_ and 1/*γ’* _*P*_. The proportion of asymptomatic infection was defined as *d*_*P*_. The *S*_*P*_ will be infected through sufficient contact with *W* and *I*_*P*_, and the transmission rates were defined as *β*_*W*_ and *β*_*P*_, respectively. We also assumed that the transmissibility of *A*_*P*_ was *κ* times that of *I*_*P*_, where 0 ≤ *κ* ≤ 1.

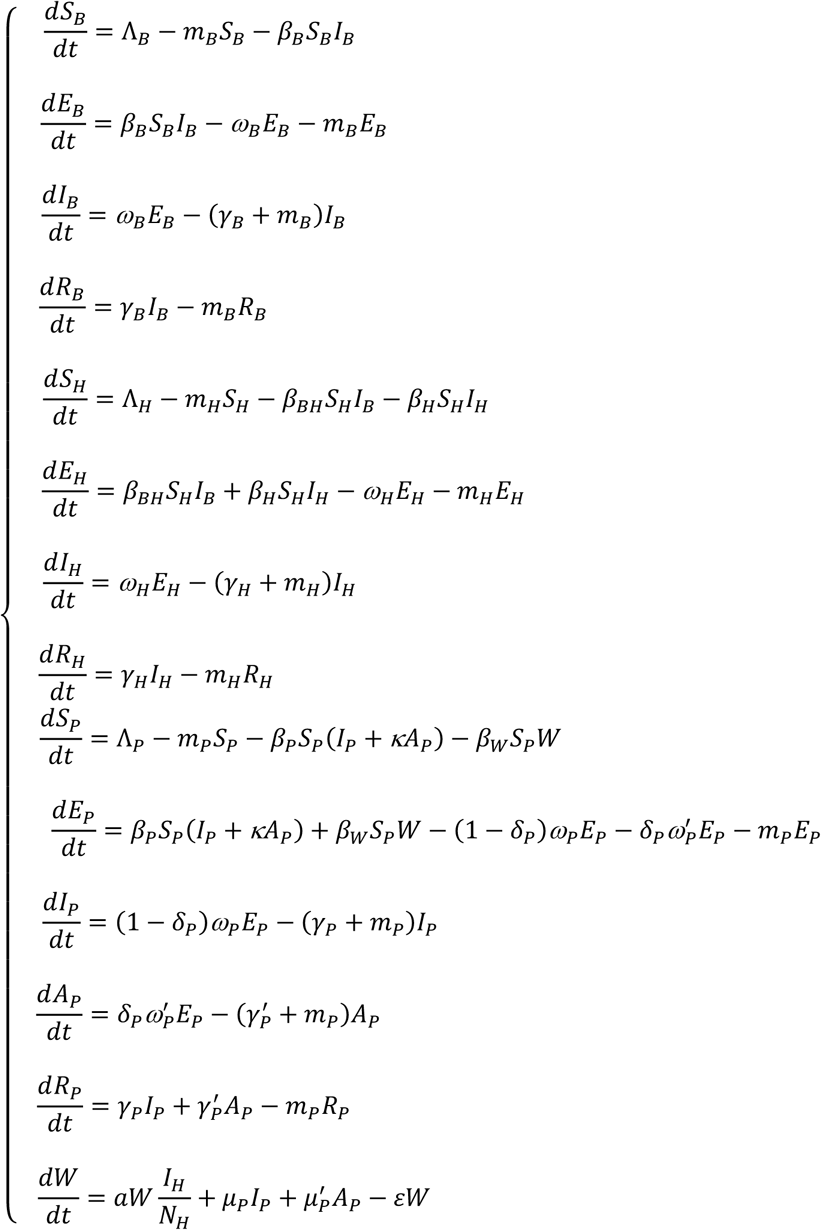

The parameters of the BHRP model were shown in Table 1.

**Table 1.**
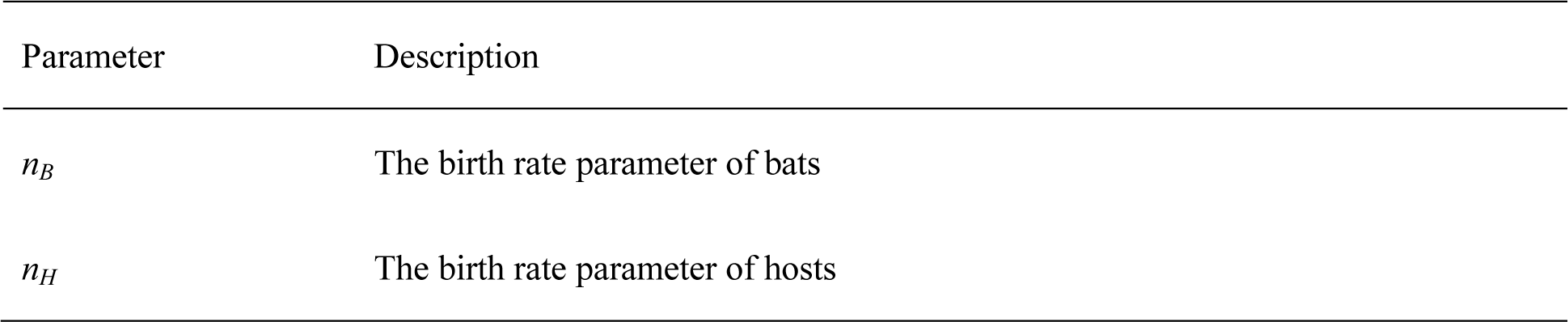
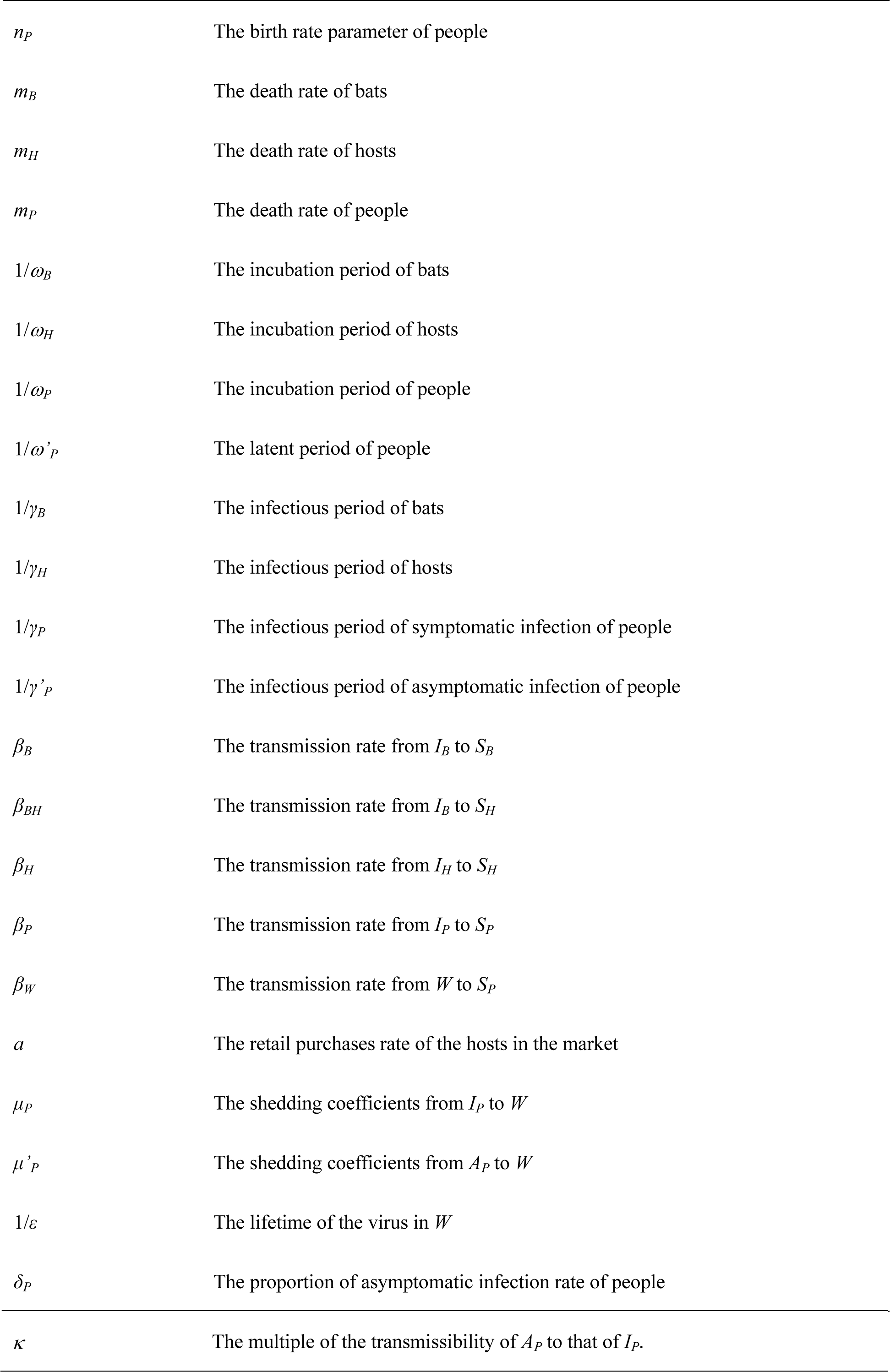
Definition of those parameters in the BHRP model.

### The simplified Reservoir-People transmission network model

Based on the information we known, we assumed that the 2019-nCoV might be imported to the seafood market in a short time. Therefore, we added the further assumptions as follows:

a. The transmission network of Bats-Host was ignored.
b. Based on our previous studies on simulating importation(*3, 4*), we set the initial value of *W* as following impulse function:

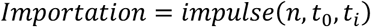

In the function, *n, t*_0_ and *t*_*i*_ refer to imported volume of the 2019-nCoV to the market, start time of the simulation, and the interval of the importation.

Therefore, the BHRP model was simplified as RP model and is shown as follows:

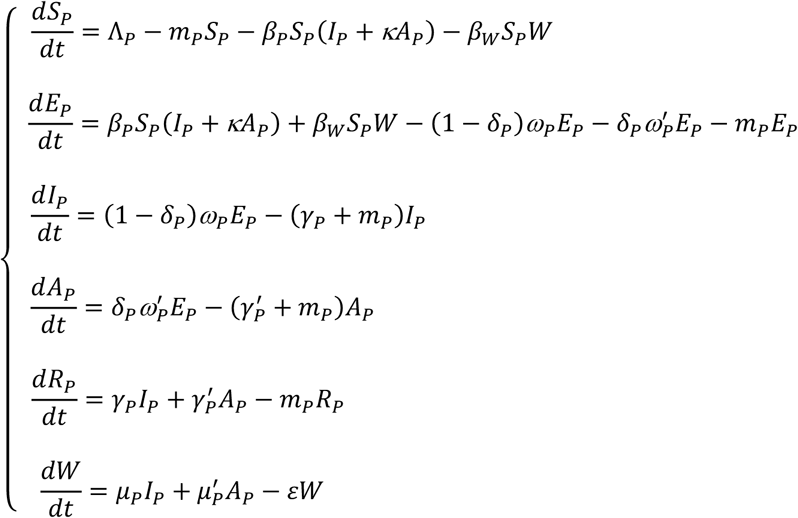

During the outbreak period, the natural birth rate and death rate in the population was in a relative low level. However, people would commonly travel into and out from Wuhan City due to the Chinese New Year. Therefore, *n*_*P*_ and *m*_*P*_ refer to the rate of people traveling into Wuhan City and traveling out from Wuhan City, respectively.

### The transmissibility of the 2019-nCoV based on the RP model

In this study, we used the basic reproduction number (*R*_0_) to assess the transmissibility of the 2019-nCoV. Commonly, *R*_0_ was defined as the expected number of secondary infections that result from introducing a single infected individual into an otherwise susceptible population(*3*). If *R*_0_ > 1, the outbreak will occur. If *R*_0_ < 1, the outbreak will go to an end.

Based on the equations of RP model, we can get the disease free equilibrium point as:

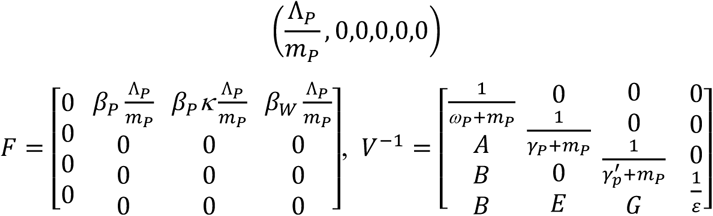

In the matrix:

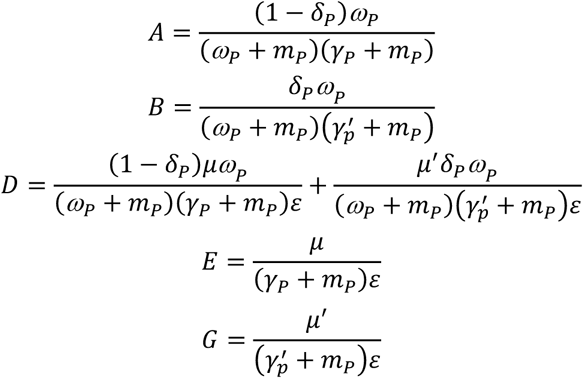

By the next generation matrix approach(*5*), we can get the next generation matrix and *R*_0_ for the RP model:

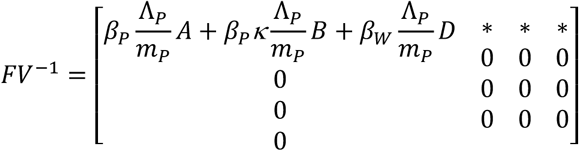

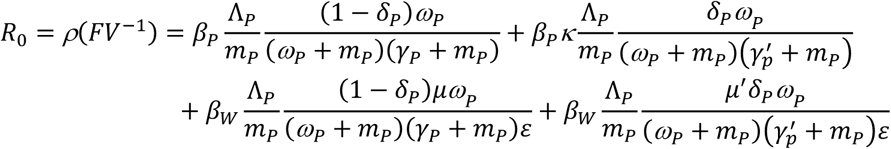

## Author contribution statements

**Tianmu Chen:** Methodology, Formal analysis, Writing - original draft, review & editing. **Jia Rui:** Methodology, Formal analysis. **Zeyu Zhao:** Formal analysis. **Qiupeng Wang:** Formal analysis. **Jing-An Cui:** Methodology. **Ling Yin:** Methodology.

## Acknowledgment

None.

